# The Coli Toolkit (CTK): An extension of the modular Yeast Toolkit for use in *E. coli*

**DOI:** 10.1101/2025.07.09.663901

**Authors:** Jacob Mejlsted, Erik Kubaczka, Sebastian Wirth, Heinz Koeppl

## Abstract

Genetic circuits are a cornerstone of synthetic biology, enabling programmable control of cellular behavior for applications in health, sustainability, and biotechnology. While Genetic Design Automation (GDA) tools have optimized and streamlined the design of such circuits, rapid and efficient assembly of DNA remains a bottleneck in the DBTL cycle. Here we present the Coli Toolkit (CTK), a modular Golden Gate-based cloning system, adapted from the Yeast Toolkit (YTK) for use in *Escherichia coli*. The CTK expands on the original YTK architecture by introducing a more flexible control of transcription and translation through subdividing the former promoter part into subparts; promoter, insulating ribozyme, and ribosome binding site (RBS). We provide a range of basic parts that enable the assembly of a wide range of constructs, as well as characterization data for all constitutive and inducible promoters provided. Additionally, we provide characterization data, as well as calibrated models, for all 20 NOT gates from the Cello library, and we provide the NOT gates as preassembled basic parts, which enables rapid cloning of larger genetic circuits. With this toolkit, we leverage the strengths of the YTK architecture to enable rapid and high-efficiency assembly of genetic circuits in *E. coli*, filling a key gap in the infrastructure of bacterial synthetic biology.

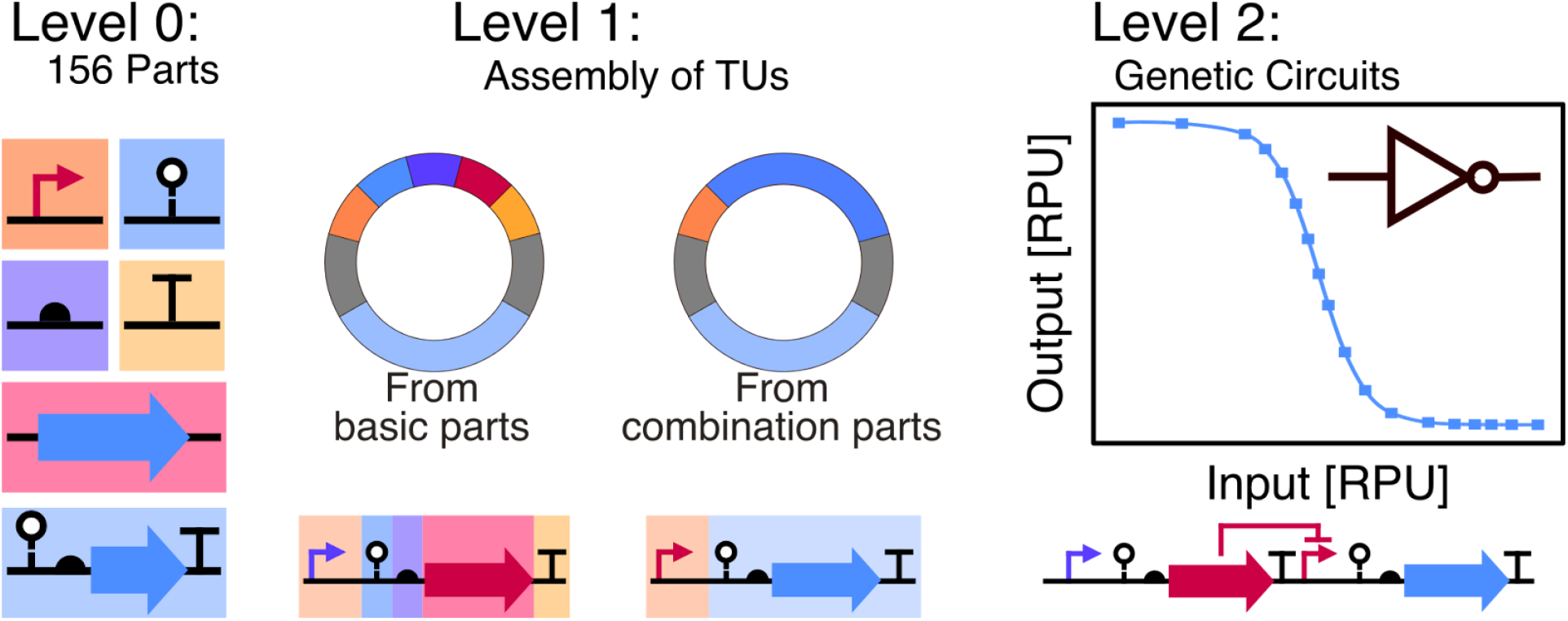

## Introduction

Synthetic biology aims to apply engineering principles and frameworks to living materials, such as organisms and cells. One aspect of this is genetic circuits, which have great potential to address many of the challenges that humankind faces, ranging from human health through diagnostics and treatments, to sustainability through adaptability and responsiveness [1], [2]. Within the design-build-test-learn (DBTL) cycle, Genetic Design Automation (GDA) tools like Cello [3], [4] and ARCTIC [5], [6], [7], have revolutionized the design stage through the determination of optimal circuit configurations based on experimental characterization data. However, the limiting step in the DBTL cycle has moved to the build stage, where the DNA encoding these circuits needs to be assembled. Standardized toolkits can enable fast, standardized and reproducible assembly of complex functions from basic parts [8]. These beneficial aspects have already been shown for other subfields within synthetic biology, like metabolic engineering, but are still lacking for genetic circuits [9].

The Gram-negative bacterium *Escherichia coli* is one of the most used organisms within synthetic biology, both for basic research, but also for various applications. It is therefore natural that multiple cloning standards already exist for it, including MoClo [8], CIDAR MoClo [10], GoldenBraid [11], [12], [13], and EcoFlex [14], among others. However, to achieve predictable circuit function, research has shown that insulating ribozymes between the promoter and RBS is essential [3]. The aforementioned cloning standards for *E. coli* does not have the ability to include an insulating ribozyme between promoter and RBS without drastically reshaping the architecture, and thereby removing backwards-compatibility. An alternative is the highly efficient and well-characterized MoClo-Yeast Toolkit (YTK) [15]. As the name states, this was originally developed for the yeast *Saccharomyces cerevisiae*, but has since been expanded by various research groups to include functions such as multiplex integration (MYT) [16], GPCR sensors [17], optogenetics (yOTK) [18], polycistronic expression [19], secretion and display [20], insulated transcriptional units for genetic circuits [Maik Molderings, unpublished results]. Additionally, the YTK has also been the basis for toolkits for other organisms such as for fission yeast, *Schizosaccharomyces pombe* (POMBOX) [21], *Kluyveromyces marxianus* [22], *Pichia pastoris* (now *Kromatogella phaffi*) [23], *Candida glabrata* [24], mammalian cells (MTK) [25], and in the bee gut microbiome (BTK) [26].

The YTK is a hierarchical cloning system in the style of MoClo. It starts with basic plasmids (level 0) that each carry a part with an abstracted biological function, such as “promoter” or “CDS”. Each basic plasmid has a part type, numbered 1-8, with optional subtypes, that determines their order. What determines the part type is the 4 base pair overhang that gets produced when the part is cut with a BsaI restriction enzyme. For example, if a part has AACG as the overhang on the 5’ end, and TATG on the 3’ end, this will be a Type 2 part. The basic parts can be assembled into cassette plasmids (level 1) that typically contain one transcriptional unit. All 8 types have to be present in order to assemble a functional cassette plasmid. However, to reduce the amount of separate DNA fragments cloned together in one reaction, combination parts can be used. For example, most cloning backbones are Type 678, which means that they have the Type 6 overhang at the 5’ end and Type 8 overhang at the 3’ end. The level 1 cassette plasmids can subsequently be combined into multigene plasmids (level 2). The order of the different transcriptional units in the multigene plasmid is determined by the assembly connectors (Type 1 and 5) from the previous level. Up to 10 transcriptional units can be assembled together using the existing assembly connector parts [15], [16].

The main advantages of the YTK over many other MoClo assembly standards for bacteria are threefold: First, the cloning overhangs in the YTK are very orthogonal to each other [27], [28] (Supplementary Figure S1). This enables the assembly of more fragments in one reaction, which, in turn, can increase customizability, without compromising the cloning efficiency. Second, the YTK contains not only well-characterized parts, but also robust support for cloning infrastructure. Connectors, origins of replication, and resistance markers exist both as individual parts, but also as combination parts to ease assemblies [15], [16]. Third, the ability to have both split parts (like Type 3a and 3b), but also combination parts greatly increase the flexibility of the system and allows for design outside the scope of the original toolkit, while maintaining backwards compatibility and the general integrity of the system.

Here, we introduce the Coli Toolkit (CTK), a toolkit for work in prokaryotes that extends the existing infrastructure of the YTK, and allows synthetic biologists to more easily work with genetic circuits in *E. coli*. To extend the capabilities of the toolkit into *E. coli*, we split the Type 2 promoter part into four subparts, which allows for independent choice of promoter(s), insulating ribozyme, and RBS. CTK comprises 156 parts, including 45 promoters, 12 ribozyme insulators, 21 RBSs, 20 CDSs, and 15 transcriptional terminators. Furthermore, we also provide 16 level 0 backbones for cloning and characterization with various combinations of origins, resistances and counter-screenable markers. Additionally, the CTK also includes 25 combination parts that enable fast and efficient cloning, of which 20 are characterized NOT gates ready to use in larger genetic logic circuits.

## Results

### Design of the Toolkit

The Coli ToolKit (CTK) is based on the YTK, with certain additions that make it possible to use it in the context of a prokaryotic host like *E. coli*. As mentioned above, the YTK works by having 8 different types of parts, numbered 1-8, that each have an abstracted function. For example, Type 1 parts are assembly connectors, Type 2 are promoters, Type 3 are coding sequences, etc. To adapt this system to making genetic circuits in *E. coli*, the Type 2 was recontextualized from only being a promoter, to transcriptional and translational control, and subsequently split into multiple sub-parts (Figure 1). Now, Type 2a and 2b are promoter parts, Type 2c is a ribozyme part and Type 2d is an RBS part. For the promoters, it is possible to use tandem promoters with one promoter in Type 2a and one in Type 2b (Figure 1A), or using a single promoter in a Type 2ab part (Figure 1B).

**Figure 1:**
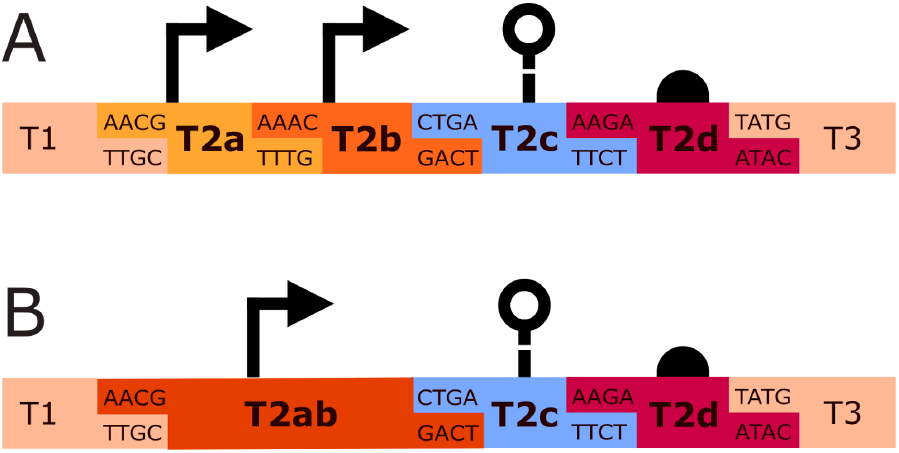
CTK Type 2 part subdivision. (A) The configuration for the use of tandem promoters with Type 2a and Type 2b parts can be seen. (B) The configuration for a single promoter as a Type 2ab part can be seen. The Type 2c and 2d parts stay consistent independently of whether a single or tandem promoters are used.

When splitting one part into multiple, suitable overhangs need to be selected to maintain high cloning efficiency. The NEB Ligase Fidelity tool [27], [28], [29] was used to select additional overhangs that don’t conflict with the existing 4-base pair overhangs employed in the YTK. The new overhangs can be seen in Figure 1 and in Supplementary Figure S1.

### Parts Overview

Access to a wide range of parts is essential to a toolkit’s utility. A total of 156 basic and combination parts are therefore included in this toolkit, and an overview of them can be seen in Figure 2 below. These basic parts include promoters, ribozyme insulators, RBSs, fluorescent proteins, transcriptional repressors, and terminators. Many parts are sourced from the Cello collection [3], and the Anderson promoter library [30] and adapted to the context of the CTK. For the promoters, there are options for both tandem promoters and single promoters. Three fluorescent proteins and two transcriptional terminators are also supplied in Type 3a and 4b, respectively. Type 3a parts enable transcriptional fusions, here tagging with fluorescent proteins in the N-terminal of the CDS in a Type 3b part. When a C-terminal tag is used instead, the tag can be placed as a Type 4a part, therefore needing terminators in Type 4b [15].

**Figure 2:**
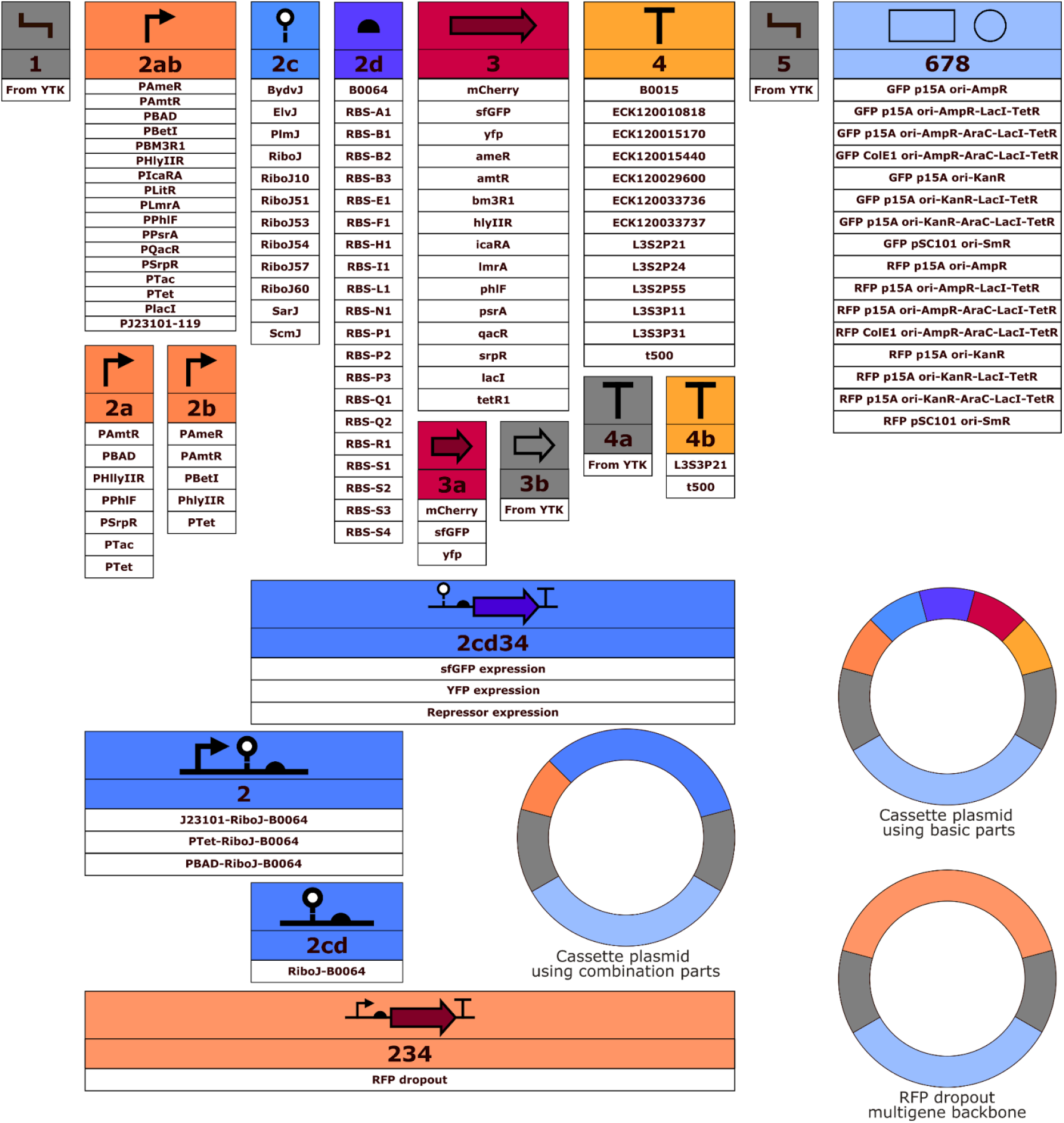
Overview of the level 0 parts in the Coli Toolkit. Each part type has unique upstream and downstream overhangs so that a full plasmid can only be assembled when all types from 1 to 8 are present. The figure shows both basic parts, and combination parts that facilitate easier cloning. The newly provided backbones are not shown in this figure. Assembly connectors (Type 1 and 5), and protein tags (Type 3b, 4a) are supplied in the YTK. A table of all plasmids, including their type, description, and resistance gene can be seen in Supplementary Table S1.

In addition to the basic parts, the CTK also contains combination parts, which are level 0 plasmids that contain multiple functional elements. These can increase the efficiency of cloning by reducing the number of parts needed for a cloning. For example, pCTK130 has the J23101 promoter (pCTK029), RiboJ ribozyme (pCTK049) and B0064 RBS (pCTK058) all in one part. Additionally, all Cello repressor units, excluding the promoter, are also included in the collection (pCTK134-pCTK153). With this, it is easier to build new genetic circuits, as only the promoter needs to be chosen. An overview of these combination parts can also be seen in Figure 2.

Several backbone plasmids of Type 678 have also been added to the toolbox. These 16 new backbones contain either an ampicillin, spectinomycin, or kanamycin resistance gene together with various origins of replication in low, medium or high copy-number. To improve on the original YTK, each cloning backbone exists in two variants; one with a GFP dropout and one with an RFP dropout. This allows for easy screening of all transcriptional units, also those that express GFP themselves. An RFP dropout (pCTK156) has also been included to enable cloning of backbone plasmids for multigene plasmids with an RFP dropout. Furthermore, some of the new backbones have repressors that regulate the input promoters (LacI regulating PTac, AraC regulating PBAD, and TetR regulating PTet). By having these regulators as part of the backbone, cloning and characterization can be streamlined as the needed regulators are already accounted for.

The full list of all plasmids can be seen in Supplementary Table S1. All part plasmids, both basic parts, combination parts, and backbone plasmids, will be submitted to Addgene for distribution during review.

### Clustering of *de novo* synthetized parts

Adding new parts to a toolkit, such as the CTK, is most commonly achieved by PCR or through *de novo* DNA synthesis. In the case of DNA synthesis, price is often a limiting factor, which means increases in efficiency are welcome.

When ordering DNA fragments from various synthesis providers, different requirements are set. During this work, the *de novo* DNA synthesis provider we ordered from required a size of the gene fragments of at least 300 bases. Smaller parts like *E. coli* promoters, RBSs, ribozymes, and terminators, are often shorter than 300 bases, even with the added overhangs for cloning into the entry plasmid, and cannot be synthesized as-is. A naïve approach would be to add random DNA at the end to pad the sequence to 300 bases before ordering, but that wastes DNA synthesis potential (Figure 3A). Instead, if multiple smaller sequences are concatenated, more parts can be ordered for the same price. To clone the parts into the entry vector without ambiguity, we exchange the BsmBI cut site with other type IIs restriction enzymes (while maintaining the overhangs for cloning into the entry vector). We can thereby package multiple fragments into one synthesis order, thus saving money and synthesis power (Figure 3B).

**Figure 3:**
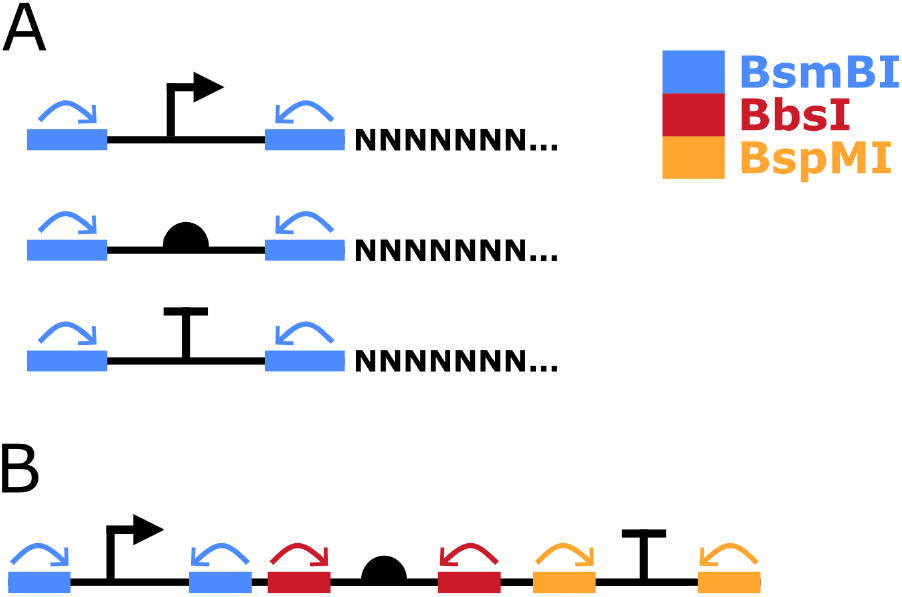
Clustering of de novo synthesized parts to increase efficiency. (A) Three individual small DNA parts can be ordered through a synthesis provider by adding filler nucleotides to obtain the minimal length required by the provider. The arrows indicate the cutting direction of the restriction enzyme. (B) Three smaller parts can be concatenated together, and have their unloading restriction enzyme altered to minimize the number of nucleotides that need to be synthesized. The arrows indicate the cutting direction of the restriction enzymes.

However, simply concatenating parts together can cause problems during synthesis due to repeats. To avoid this, one can purposefully group the parts together to avoid similarities, like concatenating a promoter together with a RBS and a terminator instead of two other promoters.

To streamline and optimize ordering *de novo* synthetized parts, we have developed a software package. It can take in DNA sequences (with the needed entry adaptors) and group them together in a way that avoids similarities, which decreases costs and increases the chances of successful DNA synthesis reactions. Through our benchmarking, we were able to decrease the costs associated with synthetizing all CTK parts to 46% of the naïve approach. (Supplementary Table S5). The software can be accessed on GitHub (See Code and Data Availability), and a benchmark comparing the software to the naïve approach and to random clustering can be seen in the Supplementary Text.

### Characterization of toolkit promoters

Constitutive promoters, like the Anderson collection [30], offer a constant level of expression that can be useful for applications. In the CTK, the Anderson promoters are included as Type 2ab parts for easy use in constructs. All have been characterized in expression of GFP through the standardized GFP expression combination part (pCTK154). The constitutive promoters have expression strengths stretching across more than two orders of magnitude (Figure 4A), and their relative strengths align with previous characterized performance from literature [30] (Supplementary Table S3). The expression from all constitutive promoters was also tested for the addition of chemical inducers used in this study, and we found that all showed unaffected behavior (Figure 4B).

**Figure 4:**
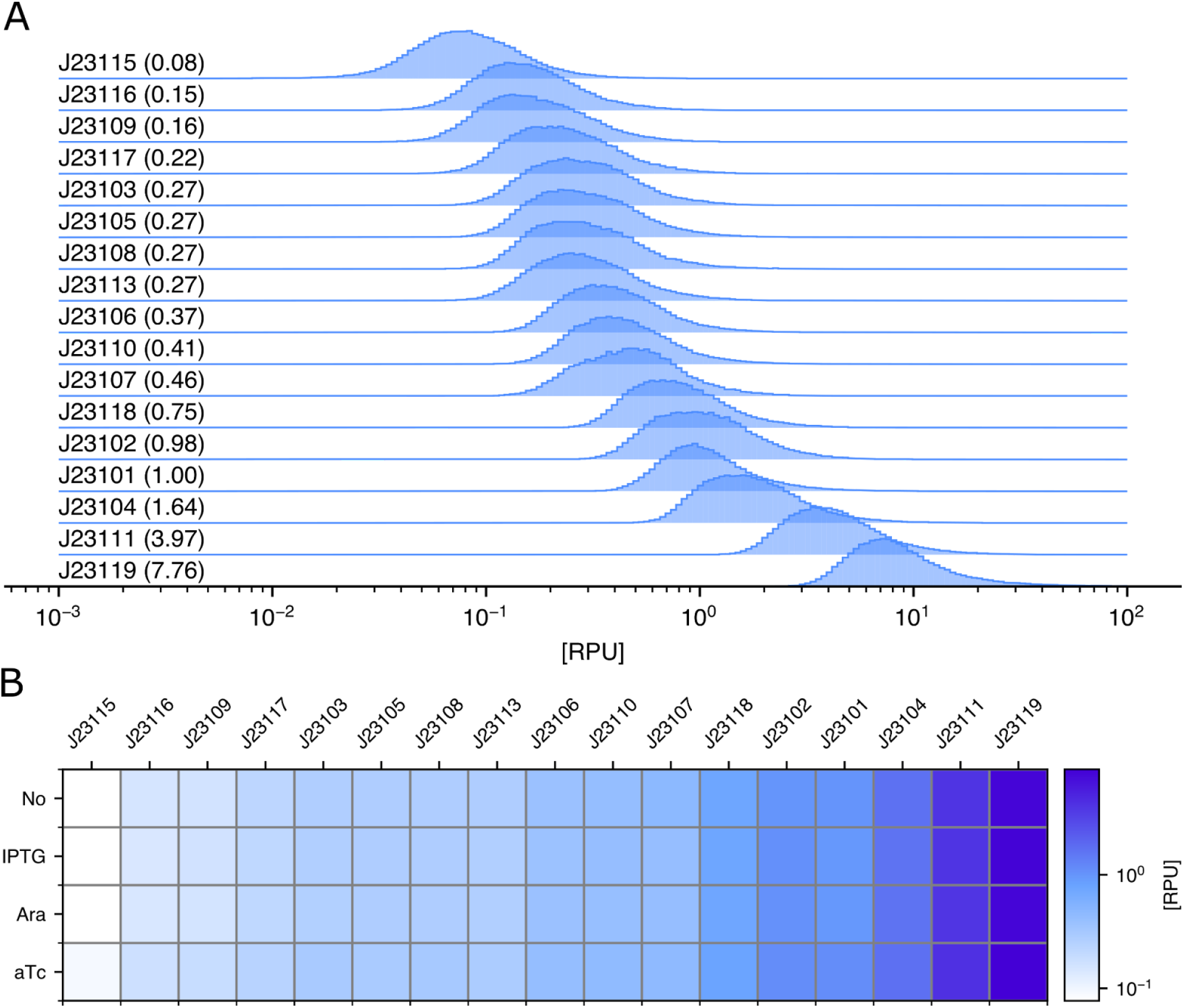
Characterization of constitutive promoters. (A) The strength of all Anderson promoters, normalized to Relative Promoter Units (RPU). Numbers in parentheses note the median expression strength of the constitutive promoter in RPU. (B) Promoter strength of Anderson promoters across different chemical inducers, normalized to RPU.

Inducible promoters allow for controlled expression based on the addition or removal of external stimuli. The most common stimulus is the addition of chemical inducers [31], but there are many other options like light [18], temperature [32], and magnetic fields [33]. To construct inputs for the genetic circuits, we are employing three promoters which respond to chemicals; PTac (responds to IPTG), PTet (responds to anhydrotetracycline (aTc)), and PBAD (responds to arabinose). These inducible promotes were sourced from the Cello collection [3].

The inducible input promoters were tested both with their respective chemical inducers (Figure 5A), but also the other inducers to measure their orthogonality (Figure 5B). From this, we can see that all input promoters act orthogonally depending on the input of chemical inducers.

**Figure 5:**
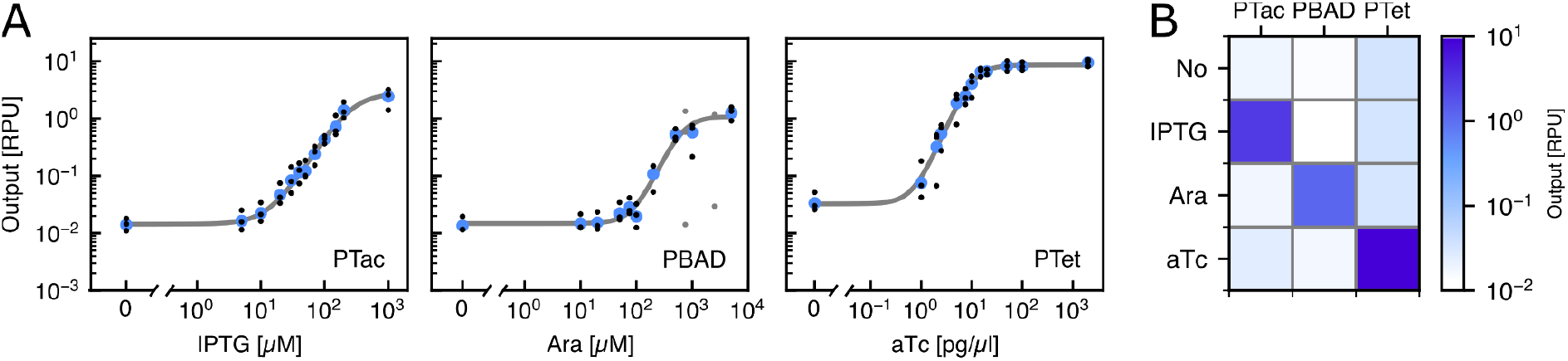
Characterization of inducible promoters. (A) The dose-response curve of the three inducible promoters in Relative Promoter Units (RPU). PTac is induced by IPTG (left), PBAD is induced by L-arabinose (middle), and PTet is induced by aTc (right). Blue points are medians of the three replicates in black. Grey dots are outliers that were not considered further (see Methods). Population distributions for all input measurements can be seen in Supplementary Figure S2. Parameters for dose-response curves can be seen in Supplementary Table S4. (B) Cross reactivity of inducible promoters. The inducible promoters only activate expression when exposed to their cognate chemical inducer. Otherwise, the expression is low.

### Characterization of Cello NOT gates

The basic components of many genetic circuits are NOT and NOR gates based on protein repressors. In the Cello library, there are a total of 20 NOT gates that can be combined in various ways to create the desired genetic circuits. For the GDA software to work, detailed characterization is required for all individual NOT gates.

All 20 NOT gates were assembled from the basic parts in the CTK, and subsequently characterized by flow cytometry. Each gate consists of the Ptac promoter driving the expression of the protein repressor in the first transcriptional unit. In the second transcriptional unit, the corresponding promoter is driving the expression of sfGFP (Figure 6A). For each gate, 12 different concentrations of IPTG were used for characterization (Figure 6B), from which a Hill curve was fit (Figure 6C). The fit takes into account the spread and density of the measured populations, which provides a more accurate response function. The parameters of the response functions of all NOT gates can be seen in Supplementary Table S5, and the individual response functions for each NOT gate can be seen in Supplementary Figure S4. The five NOT gates with highest dynamic range are shown in Figure 6D to highlight their interoperability in future genetic circuits.

**Figure 6:**
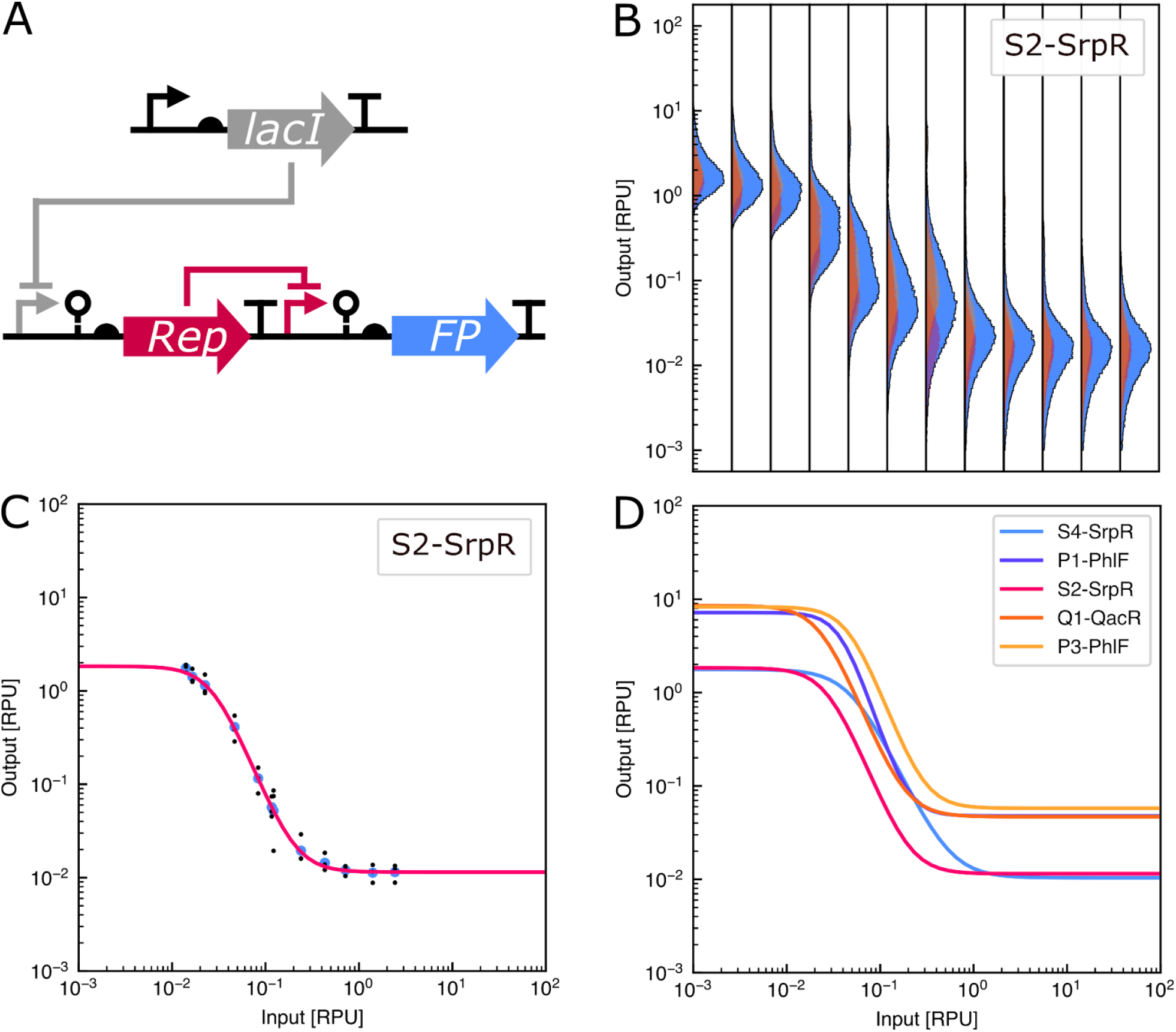
Characterization of NOT gates in the CTK. (A) The structure of the NOT gates have lacI expressed from the plasmid backbone with the NOT gate itself being composed of two transcriptional units. The first is the Repressor protein being expressed from the PTac promoter. In the second transcriptional unit, the corresponding promoter drives the expression of GFP. (B) Individual populations of the S2-SrpR across 12 different concentrations of IPTG with increasing concentrations going to the right. The individual replicates can be seen in yellow, orange and pink. The larger peak in light blue is the merged population of the replicates. Output normalized to RPU. (C) Fitting of a Hill-curve to the median of the merged three biological replicates from separate days. Black points are medians for the individual replicates, blue points are the medians of the merged populations of all replicates. (D) Fitted Hill-curves of the five NOT gates with the highest dynamic range, normalized to input in RPU and output in RPU. Cell populations as measured by flow cytometry for all 20 NOT gates can be found in Supplementary Figure S3, and fitted curves for all 20 NOT gates can be found in Supplementary Figure S4.

## Conclusion

In this paper, we have presented an expansion of the Yeast Toolkit to the prokaryote *E. coli*. To adapt the Yeast Toolkit to *E. coli*, we have expanded the scope of the type 2 parts from being just promoters to controlling transcription and translation. By splitting type 2 parts into 4 subtypes, promoters, ribozymes and RBSs can be chosen independently, thus expanding the scope of possible constructs. A toolkit also needs useful parts, so we have characterized a collection of constitutive promoters, ranging two orders of magnitude, and a set of chemically inducible input promoters that all show low leakage, a high dynamic range, and no cross-talk between them. Additionally, we characterized all 20 NOT gates from the Cello collection in the context of the CTK. The NOT gates are also provided as combination parts with ribozyme, RBS, coding sequence and terminator in one part, to facilitate faster cloning of larger genetic circuits. Lastly, we also provide a software package to increase the efficiency of *de novo* DNA synthesis orders. By using the clustering package, smaller DNA fragments can be grouped together to not waste synthesis potential. This software tool can help users of the CTK, and any other Golden Gate toolkit, to greatly decrease their synthesis costs of small DNA parts.

In this work, we have applied the CTK to genetic circuits, but it is not limited to only one topic within synthetic biology. Both applied and basic research is being conducted within our research group in *E. coli* using the CTK. We therefore believe all of these contributions will be useful to the wider synthetic biology community, both for making and using genetic circuits, but also for projects where the modular structure of the CTK can be used to easily and efficiently clone the desired constructs.

## Supporting information

CTK Plasmids

## Author Contributions

J.M. conceptualized the project, performed all experiments, and wrote the manuscript draft.

E.K. provided the flow cytometry analysis, developed the model calibration pipeline and contributed to figure creation.

S.W. implemented the clustering of *de novo* synthesized parts.

H.K. conceptualized and supervised the project.

All authors revised the manuscript.

## Acknowledgments

The authors thank Maik Molderings for lending the L1-LEU vectors, Vincent Gunawan from the Süß lab for assistance with the flow cytometry measurements, and to Anika Kofod Petersen for her work on the prototype of the *de novo* synthesis clustering pipeline. The work was made possible with the support of a scholarship from the German Academic Exchange Service (DAAD), project number 91877921 to J.M. E.K. was supported by ERC-PoC grant PLATE (101082333). Any opinions, findings, and conclusions or recommendations expressed in this material are those of the author(s) and do not necessarily reflect the views of the funding agencies.

## Materials and Methods

### Strains and growth media

Characterization experiments were performed in DH10B cells (NEB). DH10B and TOP10 cells (ThermoScientific) were used for routine transformations and cloning of plasmids. Routine bacterial growth was performed in LB media (Carl Roth), and characterization experiments were performed in Hi-Def Azure Media (Teknova), supplemented with 1% glucose. All cells were made chemically competent using the Mix & Go! kit (Zymo Research). For antibiotic selection, the following concentrations were used: Ampicillin (Amp, 100 µg/ml), kanamycin (Kan, 50 µg/ml), chloramphenicol (Cm, 25 µg/ml), spectinomycin (Spec, 50 µg/ml).

### Cloning of CTK parts

Creation of level 0 CTK parts were created in three ways: Most parts were synthesized *de novo* by Twist Biosciences (Twist Bioscience) with appropriate overhangs. For the smaller of the parts, multiple were concatenated within one linear fragment, and the BsmBI cut sites were exchanged with BbsI and BspMI to allow for targeted entry into pre-digested pYTK001, as described above. Additionally, promoter PlacI was created through oligo annealing, where overhangs were already present, and the fragment could be cloned directly into pYTK001 [15]. Construction of the Type 678 cloning backbones was performed by PCR amplification from pAN1717 and pAN3938 [3], using a Q5 polymerase (NEB), and was combined by Hi-Fi assembly (NEB) with the BsmBI overhangs and GFP expression unit from pYTK001, using the manufacturers instructions. pAN1717 and pAN3938 were gifts from Christopher Voigt (Addgene plasmid #74696 and 74697, respectively).

Cloning of level 1 and level 2 plasmids was performed using as described in the original YTK paper [15]. For assemblies with more than 4 or more parts, and for assemblies that exhibited low efficiencies, inspiration was taken from the CIDAR MoClo protocol [10], where reactions were incubated in a thermocycler for 30-60 cycles of 37 or 42 °C (5 min), depending on whether the assembly uses BsaI or BsmBI, respectively, and 16 °C (5 min), followed by final digestion (37 or 55 °C, 20 min) and enzyme inactivation (80 °C, 10 min). Constructs were checked by colony PCR using DreamTaq PCR (ThermoScientific), and sequence verified by Sanger sequencing (Microsynth) and Nanopore sequencing (Microsynth).

### Clustering algorithm for *de novo* synthesis of DNA parts

The clustering software uses the Levenshtein similarity matrix to compute the differences between the various fragments that the user wants to synthesize. Using affinity propagation [34], the software defines clusters with high sequence similarity. From this, groups are made of up to three sequences from distinct clusters to obtain low sequence similarity in the final DNA sequence sent for synthesis. If the aggressive clustering option is selected, groups only containing one sequence are concatenated together to minimize the amount of DNA needed to be synthesized. Following the grouping, the DNA sequences are concatenated and the restriction sites for BsmBI are exchanged to BbsI and BspMI for the second and third occurrences, respectively. The final sequence is then outputted as a .csv file to the same folder as the input file was chosen from.

Benchmarking was performed by running the sets of basic parts through the clustering software, both on aggressive clustering and normal clustering. The random and naive groupings were generated by a modified version of the software that bypasses the clustering. All outputs were uploaded to the website of Twist Biosciences and the ability to synthesize and price was recorded.

### Flow cytometry measurements of GFP expression

To measure the fluorescence of the individual cells, colonies were added to 200 µl Hi-Def Azure Media (Teknova), supplemented with 1% glucose and appropriate antibiotics in a 96 DeepWell plate (ThermoScientific) and grown overnight at 37 °C and 1000 RPM shaking. The culture was diluted twice by mixing 15 µl culture to 185 µl media, to a total dilution of 1:152. To this, inducers were added, and the cultures were incubated at 37 °C for 5 hours at 1000 RPM.

The used inducers were IPTG (ThermoScientific), L-arabinose (Carl Roth), and anhydrotetracycline (aTc) (ThermoScientific). For IPTG, the following final concentrations were used: 0, 5, 10, 20, 30, 40, 50, 70, 100, 150, 200, and 1000 μM.

For L-arabinose, the following final concentrations were used: 0, 10, 20, 50, 75, 100, 200, 500, 750, 1000, 2500, and 5000 µM. For aTc, the following final concentrations were used: 0, 1, 2, 2.5, 5, 7.5, 10, 15, 20, 50, 100, and 2000 pg/µl. Flow cytometry was performed on a CytoFLEX S cytometer (Beckman Coulter). For the measurements, the culture was diluted 1:20 by adding 10 µl culture to 190 µl PBS. The flow cytometer was set to record 3 . 10^5^cells for all samples.

### Flow cytometry data preprocessing and cleaning

Following acquisition of flow cytometry data, gating and data analysis was performed using a Python script (See Code and Data Availability). In particular, cytometry data was gated with the FlowCal library [35] by density gating defined on forward scatter area and height to preserve 95% of the cell events. For data cleaning, we consider the pair-wise ratios of the replicates’ median values and discard the data for the considered experimental setting (all three replicates for single inducer concentration) if not all ratios are smaller or equal to eight, while otherwise the samples are pooled together. In addition, we only consider replicates with at least 1000 cell events. Both procedures were chosen to ensure robust data cleaning.

Further preprocessing includes conversion to relative promoter units (RPU) to quantify the relative promoter activity in comparison to the reference promoter J23101 [3], [36]. The conversion factor γ to convert raw fluorescence intensity values into RPU was determined by using the following formula from [3]:

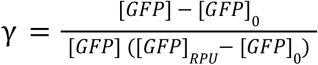

Here, [*GFP*] is the median fluorescence of the sample, [*GFP*]_0_ is the median fluorescence of the autofluorescence of the control, and [*GFP*]_*RPU*_ is the median fluorescence of the cells containing the reference plasmid with the J23101 promoter [3], [36]. All raw values then have been rescaled by multiplication with γ to yield RPU values.

### Model calibration

To represent the dose-response curves of input sensors and gates analytically, we calibrate models to the median RPU dose-response. In particular, this means that model calibration uses the dataset *D* = {*x*_*i*_, *y*_*i*_}. Here, the *x*_*i*_ ∈ ℝ_>0_ either represent the inducer concentrations in case of the input sensors or the corresponding input sensor’s median output in RPU, representing the gate’s input, for gate calibration. In both cases the *y*_*i*_ ∈ ℝ_>0_ are the corresponding median outputs in RPU.

As models, we use the activatory Hill equation defined as

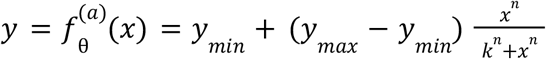

for the input sensors and the inhibitory hill equation

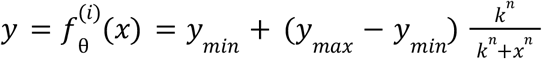

for the gates. Parameters *y* _*min*_, *y* _*max*_, and *k* are in RPU and define the dynamic range of the output (*y* _*min*_ and *y* _*max*_) as well as the location of the transition region (*k*). *n* is the Hill coefficient and defines the steepness, respectively the length of the transition region. θ = (*y* _*min*_, *y* _*max*_, *n, k*) represents the model’s parameterization.

The algorithm for model calibration we use is parallel tempering [37], [38], a Markov chain Monte Carlo algorithm. Parallel tempering uses Markov chains at different temperatures to draw samples from the posterior distribution, allowing to explore multimodal distributions through sample exchange between chains, and was successfully applied to model calibration of chemical reaction networks previously [39]. As the experimental values as well as the parameters θ span multiple orders of magnitude, we will consider in both cases the logarithmic domain for the calculation of differences. For the experimental values this ensures that the model’s deviation to the data is treated in dependence to the order of magnitude, while in case of the parameters it improves the algorithmic behavior.

The posterior distribution p(θ|*D*) over the parameters θ in dependence to the median dose-response data *D* is defined in terms of the prior p(θ) and the likelihood p(*D*|θ) as

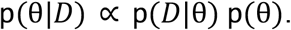

The prior encodes our initial assumptions on the parameters. We define the prior to be

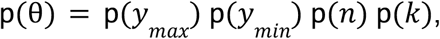

and assume that for *y* _*max*_ and *y* _*min*_ values in the range [ŷ _*max*_, 2 ŷ _*max*_] and [0. 5 ŷ _*max*_, ŷ _*min*_] respectively should be most likely, where 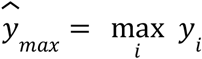 and 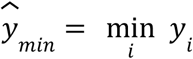. We encode this as

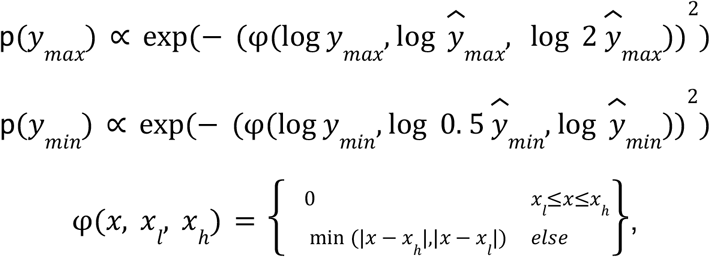

while we set p(*n*) = p(*k*) = 1, encoding no further assumptions. Please note that defining the distributions in terms of proportionalities only is sufficient for maximum *a posteriori* estimation and sampling from the posterior by parallel tempering.

The likelihood p(*D*|θ) characterizes how well the data matches the model with parameters θ. As both models (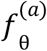 and 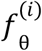) have four parameters each, they can be treated identically wherefore we introduce *f*_θ_ as a placeholder representing either of 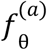 and 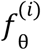. We assume independence of data points wherefore we define the factorized likelihood

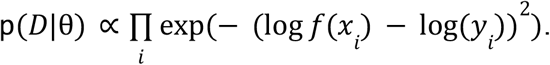

The calibrated parameter configuration 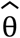 is then defined as the maximum *a posteriori* (MAP) estimate

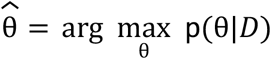

derived by sampling from the posterior distribution with parallel tempering. In particular, we define the initial parameter configuration to be

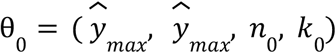

With *n*_0_ = 2 and *k*_0_ = 1 in the case of input sensor calibration or *k*_0_ = 0. 01 in the case of gate calibration. Then, we executed parallel tempering with 10 walkers, each featuring 10 chains at different temperatures, for 10,000 steps. From the 10^*7*^ posterior evaluations we select 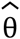 as the one maximizing it. This process is done independently for each input sensor and each gate.

The full model calibration pipeline is part of the Python script for data processing (Code and Data Availability).

## Code and Data Availability

The data and code is available at

https://github.com/Self-Organizing-Systems-TU-Darmstadt/CTK-ColiToolKit

## Plasmid and Sequence Availability

All CTK plasmid sequences are available from the attached zip file and will be made available through Addgene during the review. Additional cloned plasmids are available upon request.

## Supplementary Information

In the supplementary, tables showing the 156 CTK plasmids provided in the toolkit; the plasmids cloned and used for characterization experiments; characterization data for constitutive promoters, inducible promoters, and NOT gates. Figures describing the new CTK overhangs for the type 2 subparts; flow cytometry distributions of input promoters and NOT gates, and hill curve fits for all NOT gates. Additionally, supplementary text describing the benchmarking of the clustering algorithm. Lastly, sequence files in GenBank format for the 156 CTK plasmids included in the toolkit.

