## Supplementary information for "The Coli Toolkit (CTK): An extension of the modular Yeast Toolkit for use in *E. coli*"

### Supporting Tables and Figures

| | |
|---|---|
| Supplementary Table S1 | List of Coli Toolkit plasmids |
| Supplementary Table S2 | List of plasmids made for characterization results |
| Supplementary Table S3 | Constitutive promoter expression levels |
| Supplementary Table S4 | Inducible promoters dose-response parameters |
| Supplementary Table S5 | NOT gate response function parameters |
| Supplementary Figure S1 | Ligase fidelity overview of the 4-base pair overhangs employed in the CTK |
| Supplementary Figure S2 | Distributions of input promoters |
| Supplementary Figure S3 | Distributions of NOT gates |
| Supplementary Figure S4 | Response functions of NOT gates. |

### Supporting Text

| | |
|---|---|
| Supplementary Text | Benchmarking clustering algorithm |
| Supplementary Table S5 | Benchmark of set 1, the CTK parts |
| Supplementary Table S6 | Benchmark of set 2, the parts required for the 2-bit hashing function |
| Supplementary Table S7 | Benchmark of set 3, RcsAB promoters |

### Supplementary Table S1: List of Coli Toolkit plasmids

| *Plasmid* | *Type* | *Description* | *Resistance* |
|---|---|---|---|
| pCTK001 | 2ab | PAmeR | CamR |
| pCTK002 | 2ab | PAmtR | CamR |
| pCTK003 | 2ab | PBAD | CamR |
| pCTK004 | 2ab | PBetI | CamR |
| pCTK005 | 2ab | PBM3R1 | CamR |
| pCTK006 | 2ab | PHlyIIR | CamR |
| pCTK007 | 2ab | PIcaRA | CamR |
| pCTK008 | 2ab | PLitR | CamR |
| pCTK009 | 2ab | PLmrA | CamR |
| pCTK010 | 2ab | PPhlF | CamR |
| pCTK011 | 2ab | PPsrA | CamR |
| pCTK012 | 2ab | PQacR | CamR |
| pCTK013 | 2ab | PSrpR | CamR |
| pCTK014 | 2ab | PTac | CamR |
| pCTK015 | 2ab | PTet | CamR |
| pCTK016 | 2ab | PLacI | CamR |
| pCTK017 | 2a | PAmtR | CamR |
| pCTK018 | 2a | PBAD | CamR |
| pCTK019 | 2a | PHlyIIR | CamR |
| pCTK020 | 2a | PPhlF | CamR |
| pCTK021 | 2a | PSrpR | CamR |
| pCTK022 | 2a | PTac | CamR |
| pCTK023 | 2a | PTet | CamR |
| pCTK024 | 2b | PAmeR | CamR |
| pCTK025 | 2b | PAmtR | CamR |
| pCTK026 | 2b | PBetI | CamR |
| pCTK027 | 2b | PhlyIIR | CamR |
| pCTK028 | 2b | PTet | CamR |
| pCTK029 | 2ab | J23101 | CamR |
| pCTK030 | 2ab | J23102 | CamR |
| pCTK031 | 2ab | J23103 | CamR |
| pCTK032 | 2ab | J23104 | CamR |
| pCTK033 | 2ab | J23105 | CamR |
| pCTK034 | 2ab | J23106 | CamR |
| pCTK035 | 2ab | J23107 | CamR |
| pCTK036 | 2ab | J23108 | CamR |
| pCTK037 | 2ab | J23109 | CamR |
| pCTK038 | 2ab | J23110 | CamR |
| pCTK039 | 2ab | J23111 | CamR |
| pCTK040 | 2ab | J23113 | CamR |
| pCTK041 | 2ab | J23115 | CamR |
| pCTK042 | 2ab | J23116 | CamR |
| pCTK043 | 2ab | J23117 | CamR |
| pCTK044 | 2ab | J23118 | CamR |
| pCTK045 | 2ab | J23119 | CamR |
| pCTK046 | 2c | BydvJ | CamR |
| pCTK047 | 2c | ElvJ | CamR |
| pCTK048 | 2c | PlmJ | CamR |
| pCTK049 | 2c | RiboJ | CamR |
| pCTK050 | 2c | RiboJ10 | CamR |
| pCTK051 | 2c | RiboJ51 | CamR |

| | | | |
|---|---|---|---|
| pCTK052 | 2c | RiboJ53 | CamR |
| pCTK053 | 2c | RiboJ54 | CamR |
| pCTK054 | 2c | RiboJ57 | CamR |
| pCTK055 | 2c | RiboJ60 | CamR |
| pCTK056 | 2c | SarJ | CamR |
| pCTK057 | 2c | ScmJ | CamR |
| pCTK058 | 2d | B0064 | CamR |
| pCTK059 | 2d | RBS-A1 | CamR |
| pCTK060 | 2d | RBS-B1 | CamR |
| pCTK061 | 2d | RBS-B2 | CamR |
| pCTK062 | 2d | RBS-B3 | CamR |
| pCTK063 | 2d | RBS-E1 | CamR |
| pCTK064 | 2d | RBS-F1 | CamR |
| pCTK065 | 2d | RBS-H1 | CamR |
| pCTK066 | 2d | RBS-I1 | CamR |
| pCTK067 | 2d | RBS-L1 | CamR |
| pCTK068 | 2d | RBS-N1 | CamR |
| pCTK069 | 2d | RBS-P1 | CamR |
| pCTK070 | 2d | RBS-P2 | CamR |
| pCTK071 | 2d | RBS-P3 | CamR |
| pCTK072 | 2d | RBS-Q1 | CamR |
| pCTK073 | 2d | RBS-Q2 | CamR |
| pCTK074 | 2d | RBS-R1 | CamR |
| pCTK075 | 2d | RBS-S1 | CamR |
| pCTK076 | 2d | RBS-S2 | CamR |
| pCTK077 | 2d | RBS-S3 | CamR |
| pCTK078 | 2d | RBS-S4 | CamR |
| pCTK079 | 3 | mCherry | CamR |
| pCTK080 | 3 | sfGFP | CamR |
| pCTK081 | 3 | yfp | CamR |
| pCTK082 | 3 | ameR | CamR |
| pCTK083 | 3 | amtR | CamR |
| pCTK084 | 3 | betI | CamR |
| pCTK085 | 3 | bm3R1 | CamR |
| pCTK086 | 3 | hlyIIR | CamR |
| pCTK087 | 3 | icaRA | CamR |
| pCTK088 | 3 | litR | CamR |
| pCTK089 | 3 | lmrR | CamR |
| pCTK090 | 3 | phlF | CamR |
| pCTK091 | 3 | psrA | CamR |
| pCTK092 | 3 | qacR | CamR |
| pCTK093 | 3 | srpR | CamR |
| pCTK094 | 3 | lacI | CamR |
| pCTK095 | 3 | TetR | CamR |
| pCTK096 | 3a | mCherry | CamR |
| pCTK097 | 3a | sfGFP | CamR |
| pCTK098 | 3a | yfp | CamR |
| pCTK099 | 4 | BBa_B0015 | CamR |
| pCTK100 | 4 | ECK120010818 | CamR |
| pCTK101 | 4 | ECK120015170 | CamR |
| pCTK102 | 4 | ECK120015440 | CamR |
| pCTK103 | 4 | ECK120029600 | CamR |
| pCTK104 | 4 | ECK120033736 | CamR |
| pCTK105 | 4 | ECK120033737 | CamR |

| | | | |
|---|---|---|---|
| pCTK106 | 4 | L3S2P21 | CamR |
| pCTK107 | 4 | L3S2P24 | CamR |
| pCTK108 | 4 | L3S2P55 | CamR |
| pCTK109 | 4 | L3S3P11 | CamR |
| pCTK110 | 4 | L3S3P31 | CamR |
| pCTK111 | 4 | t500 | CamR |
| pCTK112 | 4b | L3S2P21 | CamR |
| pCTK113 | 4b | t500 | CamR |
| pCTK114 | 678 | GFP p15A ori-AmpR | AmpR |
| pCTK115 | 678 | GFP p15A ori-AmpR-LacI-TetR | AmpR |
| pCTK116 | 678 | GFP p15A ori-AmpR-AraC-LacI-TetR | AmpR |
| pCTK117 | 678 | GFP ColE1 ori-AmpR-AraC-LacI-TetR | AmpR |
| pCTK118 | 678 | GFP p15A ori-KanR | KanR |
| pCTK119 | 678 | GFP p15A ori-KanR-LacI-TetR | KanR |
| pCTK120 | 678 | GFP p15A ori-KanR-AraC-LacI-TetR | KanR |
| pCTK121 | 678 | GFP pSC101 ori - SmR | SpecR |
| pCTK122 | 678 | RFP p15A ori-AmpR | AmpR |
| pCTK123 | 678 | RFP p15A ori-AmpR-LacI-TetR | AmpR |
| pCTK124 | 678 | RFP p15A ori-AmpR-AraC-LacI-TetR | AmpR |
| pCTK125 | 678 | RFP ColE1 ori-AmpR-AraC-LacI-TetR | AmpR |
| pCTK126 | 678 | RFP p15A ori-KanR | KanR |
| pCTK127 | 678 | RFP p15A ori-KanR-LacI-TetR | KanR |
| pCTK128 | 678 | RFP p15A ori-KanR-AraC-LacI-TetR | KanR |
| pCTK129 | 678 | RFP pSC101 ori - SmR | SpecR |
| pCTK130 | 2 | J23101-RiboJ-B0064 | CamR |
| pCTK131 | 2 | PBAD-RiboJ-B0064 | CamR |
| pCTK132 | 2 | PTet-RiboJ-B0064 | CamR |
| pCTK133 | 2cd | RiboJ-B0064 | CamR |
| pCTK134 | 2cd34 | A1-amtR | CamR |
| pCTK135 | 2cd34 | B1-bm3R1 | CamR |
| pCTK136 | 2cd34 | B2-bm3R1 | CamR |
| pCTK137 | 2cd34 | B3-bm3R1 | CamR |
| pCTK138 | 2cd34 | E1-betI | CamR |
| pCTK139 | 2cd34 | F1-ameR | CamR |
| pCTK140 | 2cd34 | H1-hlyIIR | CamR |
| pCTK141 | 2cd34 | I1-IcaRA | CamR |
| pCTK142 | 2cd34 | L1-LitR | CamR |
| pCTK143 | 2cd34 | N1-LmrA | CamR |
| pCTK144 | 2cd34 | P1-phlF | CamR |
| pCTK145 | 2cd34 | P2-phlF | CamR |
| pCTK146 | 2cd34 | P3-phlF | CamR |
| pCTK147 | 2cd34 | Q1-QacR | CamR |
| pCTK148 | 2cd34 | Q2-QacR | CamR |
| pCTK149 | 2cd34 | R1-PsrA | CamR |
| pCTK150 | 2cd34 | S1-srpR | CamR |
| pCTK151 | 2cd34 | S2-srpR | CamR |
| pCTK152 | 2cd34 | S3-srpR | CamR |
| pCTK153 | 2cd34 | S4-srpR | CamR |
| pCTK154 | 2cd34 | sfGFP expression | CamR |
| pCTK155 | 2cd34 | yfp expression | CamR |
| pCTK156 | 234r | RFP dropout | CamR |

**Supplementary Table S2: List of plasmids made for characterization results**

Note: Plasmids L1-LEU-1, L1-LEU-1E, and L1-LEU-2E are backbone plasmids provided by Maik Molderings (unpublished results) that function as type 56781 parts to make cloning more efficient. These parts are very similar to the cloning backbones of the MYT kit [1] except that they contain isolating ribozymes in the backbone that isolates gene expression between transcriptional units. With the ribozymes in type 2c, this should not be required to achieve good expression results, but doesn't decrease effectiveness either.

| *Plasmid* | *Level* | *Type* | *Description* | *Resistance* | *Components* |
|---|---|---|---|---|---|
| pJCM113 | 1 | TU1 | PTac-A1-amtR | AmpR | L1-LEU-1, pCTK014, pCTK046, pCTK059, pCTK083, pCTK108 |
| pJCM114 | 1 | TU1 | PTac-B1-bm3R1 | AmpR | L1-LEU-1, pCTK014, pCTK056, pCTK060, pCTK085, pCTK108 |
| pJCM115 | 1 | TU1 | PTac-B2-bm3R1 | AmpR | L1-LEU-1, pCTK014, pCTK056, pCTK061, pCTK085, pCTK108 |
| pJCM116 | 1 | TU1 | PTac-B3-bm3R1 | AmpR | L1-LEU-1, pCTK014, pCTK056, pCTK062, pCTK085, pCTK108 |
| pJCM117 | 1 | TU1 | PTac-E1-betI | AmpR | L1-LEU-1, pCTK014, pCTK054, pCTK063, pCTK084, pCTK108 |
| pJCM118 | 1 | TU1 | PTac-F1-ameR | AmpR | L1-LEU-1, pCTK014, pCTK053, pCTK064, pCTK082, pCTK108 |
| pJCM119 | 1 | TU1 | PTac-H1-hlyIIR | AmpR | L1-LEU-1, pCTK014, pCTK051, pCTK065, pCTK086, pCTK104 |
| pJCM120 | 1 | TU1 | PTac-I1-IcaRA | AmpR | L1-LEU-1, pCTK014, pCTK047, pCTK066, pCTK087, pCTK103 |
| pJCM121 | 1 | TU1 | PTac-L1-LitR | AmpR | L1-LEU-1, pCTK014, pCTK048, pCTK067, pCTK088, pCTK106 |
| pJCM122 | 1 | TU1 | PTac-N1-LmrA | AmpR | L1-LEU-1, pCTK014, pCTK055, pCTK068, pCTK089, pCTK106 |
| pJCM123 | 1 | TU1 | PTac-P1-phlF | AmpR | L1-LEU-1, pCTK014, pCTK052, pCTK069, pCTK090, pCTK105 |
| pJCM124 | 1 | TU1 | PTac-P2-phlF | AmpR | L1-LEU-1, pCTK014, pCTK052, pCTK070, pCTK090, pCTK105 |
| pJCM125 | 1 | TU1 | PTac-P3-phlF | AmpR | L1-LEU-1, pCTK014, pCTK052, pCTK071, pCTK090, pCTK105 |
| pJCM126 | 1 | TU1 | PTac-Q1-QacR | AmpR | L1-LEU-1, pCTK014, pCTK055, pCTK072, pCTK092, pCTK105 |
| pJCM127 | 1 | TU1 | PTac-Q2-QacR | AmpR | L1-LEU-1, pCTK014, pCTK055, pCTK073, pCTK092, pCTK106 |
| pJCM128 | 1 | TU1 | PTac-R1-PsrA | AmpR | L1-LEU-1, pCTK014, pCTK057, pCTK074, pCTK091, pCTK106 |
| pJCM129 | 1 | TU1 | PTac-S1-srpR | AmpR | L1-LEU-1, pCTK014, pCTK050, pCTK075, pCTK093, pCTK103 |
| pJCM130 | 1 | TU1 | PTac-S2-srpR | AmpR | L1-LEU-1, pCTK014, pCTK050, pCTK076, pCTK093, pCTK103 |
| pJCM131 | 1 | TU1 | PTac-S3-srpR | AmpR | L1-LEU-1, pCTK014, pCTK050, pCTK077, pCTK093, pCTK103 |
| pJCM132 | 1 | TU1 | PTac-S4-srpR | AmpR | L1-LEU-1, pCTK014, pCTK050, pCTK078, pCTK093, pCTK103 |
| pJCM133 | 1 | TU2e | J23101-sfGFP | AmpR | L1-LEU-2E, pCTK029, pCTK049, pCTK058, pCTK080, pCTK106 |
| pJCM135 | 1 | TU2e | PAmeR-sfGFP | AmpR | L1-LEU-2E, pCTK001, pCTK049, pCTK058, pCTK080, pCTK106 |

| | | | | | |
|---|---|---|---|---|---|
| pJCM137 | 1 | TU2e | PAmtR-sfGFP | AmpR | L1-LEU-2E, pCTK002, pCTK049, pCTK058, pCTK080, pCTK106 |
| pJCM139 | 1 | TU2e | PBetI-sfGFP | AmpR | L1-LEU-2E, pCTK004, pCTK049, pCTK058, pCTK080, pCTK106 |
| pJCM141 | 1 | TU2e | PBM3R1-sfGFP | AmpR | L1-LEU-2E, pCTK005, pCTK049, pCTK058, pCTK080, pCTK106 |
| pJCM143 | 1 | TU2e | PHlyIIR-sfGFP | AmpR | L1-LEU-2E, pCTK006, pCTK049, pCTK058, pCTK080, pCTK106 |
| pJCM145 | 1 | TU2e | PIcaRA-sfGFP | AmpR | L1-LEU-2E, pCTK007, pCTK049, pCTK058, pCTK080, pCTK106 |
| pJCM147 | 1 | TU2e | PLitR-sfGFP | AmpR | L1-LEU-2E, pCTK008, pCTK049, pCTK058, pCTK080, pCTK106 |
| pJCM149 | 1 | TU2e | PLmrA-sfGFP | AmpR | L1-LEU-2E, pCTK009, pCTK049, pCTK058, pCTK080, pCTK106 |
| pJCM151 | 1 | TU2e | PPhlF-sfGFP | AmpR | L1-LEU-2E, pCTK010, pCTK049, pCTK058, pCTK080, pCTK106 |
| pJCM153 | 1 | TU2e | PPsrA-sfGFP | AmpR | L1-LEU-2E, pCTK011, pCTK049, pCTK058, pCTK080, pCTK106 |
| pJCM155 | 1 | TU2e | PQacR-sfGFP | AmpR | L1-LEU-2E, pCTK012, pCTK049, pCTK058, pCTK080, pCTK106 |
| pJCM157 | 1 | TU2e | SrpR-sfGFP | AmpR | L1-LEU-2E, pCTK013, pCTK049, pCTK058, pCTK080, pCTK106 |
| pJCM212 | 2 | | PTac-A1-amtR_PAmtR-sfGFP | KanR | pJCM471, pJCM113, pJCM137 |
| pJCM213 | 2 | | PTac-B1-bm3R1_PBM3R1-sfGFP | KanR | pJCM471, pJCM114, pJCM141 |
| pJCM214 | 2 | | PTac-B2-bm3R1_PBM3R1-sfGFP | KanR | pJCM471, pJCM115, pJCM141 |
| pJCM215 | 2 | | PTac-B3-bm3R1_PBM3R1-sfGFP | KanR | pJCM471, pJCM116, pJCM141 |
| pJCM216 | 2 | | PTac-E1-betI_PBetI-sfGFP | KanR | pJCM471, pJCM117, pJCM139 |
| pJCM217 | 2 | | PTac-F1-ameR_PAmeR-sfGFP | KanR | pJCM471, pJCM118, pJCM135 |
| pJCM218 | 2 | | PTac-H1-hlyIIR_PHlyIIR-sfGFP | KanR | pJCM471, pJCM119, pJCM143 |
| pJCM219 | 2 | | PTac-I1-IcaRA_PIcaRA-sfGFP | KanR | pJCM471, pJCM120, pJCM145 |
| pJCM220 | 2 | | PTac-L1-LitR_PLitR-sfGFP | KanR | pJCM471, pJCM121, pJCM147 |
| pJCM221 | 2 | | PTac-N1-LmrA_PLmrA-sfGFP | KanR | pJCM471, pJCM122, pJCM149 |
| pJCM222 | 2 | | PTac-P1-phlF_PPhIF-sfGFP | KanR | pJCM471, pJCM123, pJCM151 |
| pJCM223 | 2 | | PTac-P2-phlF_PPhIF-sfGFP | KanR | pJCM471, pJCM124, pJCM151 |
| pJCM224 | 2 | | PTac-P3-phlF_PPhIF-sfGFP | KanR | pJCM471, pJCM125, pJCM151 |

| | | | | | |
|---|---|---|---|---|---|
| pJCM225 | 2 | | PTac-Q1-QacR_PQac R-sfGFP | KanR | pJCM471, pJCM126, pJCM155 |
| pJCM226 | 2 | | PTac-Q2-QacR_PQac R-sfGFP | KanR | pJCM471, pJCM127, pJCM155 |
| pJCM227 | 2 | | PTac-R1-PsrA_PPsrA-sfGFP | KanR | pJCM471, pJCM128, pJCM153 |
| pJCM228 | 2 | | PTac-S1-srpR_PSrpR-sfGFP | KanR | pJCM471, pJCM129, pJCM157 |
| pJCM229 | 2 | | PTac-S2-srpR_PSrpR-sfGFP | KanR | pJCM471, pJCM130, pJCM157 |
| pJCM230 | 2 | | PTac-S3-srpR_PSrpR-sfGFP | KanR | pJCM471, pJCM131, pJCM157 |
| pJCM231 | 2 | | PTac-S4-srpR_PSrpR-sfGFP | KanR | pJCM471, pJCM132, pJCM157 |
| pJCM386 | 1 | TU1e | PJ23101-sfGFP | AmpR | L1-LEU-1E, pCTK029, pCTK154 |
| pJCM387 | 1 | TU1e | PTac-sfGFP | AmpR | L1-LEU-1E, pCTK014, pCTK154 |
| pJCM434 | 2 | | PJ23101-sfGFP | KanR | pJCM471, pJCM386 |
| pJCM435 | 2 | | PTac-sfGFP | KanR | pJCM471, pJCM387 |
| pJCM446 | 1 | TU1e | PBAD-sfGFP | AmpR | L1-LEU-1E, pCTK003, pCTK154 |
| pJCM447 | 1 | TU1e | PTet-sfGFP | AmpR | L1-LEU-1E, pCTK015, pCTK154 |
| pJCM448 | 2 | | PBAD-sfGFP | KanR | pJCM471, pJCM446 |
| pJCM449 | 2 | | PTet-sfGFP | KanR | pJCM471, pJCM447 |
| pJCM471 | 1 | BB | p15A ori KanR AraC-LacI-TetR | KanR | pCTK120. pYTK008, pCTK156, pYTK073 |
| pJCM534 | 1 | TU1e | PJ23102-sfGFP | AmpR | L1-LEU-1E, pCTK030, pCTK154 |
| pJCM535 | 1 | TU1e | PJ23103-sfGFP | AmpR | L1-LEU-1E, pCTK031, pCTK154 |
| pJCM536 | 1 | TU1e | PJ23104-sfGFP | AmpR | L1-LEU-1E, pCTK032, pCTK154 |
| pJCM537 | 1 | TU1e | PJ23105-sfGFP | AmpR | L1-LEU-1E, pCTK033, pCTK154 |
| pJCM538 | 1 | TU1e | PJ23106-sfGFP | AmpR | L1-LEU-1E, pCTK034, pCTK154 |
| pJCM539 | 1 | TU1e | PJ23107-sfGFP | AmpR | L1-LEU-1E, pCTK035, pCTK154 |
| pJCM540 | 1 | TU1e | PJ23108-sfGFP | AmpR | L1-LEU-1E, pCTK036, pCTK154 |
| pJCM541 | 1 | TU1e | PJ23109-sfGFP | AmpR | L1-LEU-1E, pCTK037, pCTK154 |
| pJCM542 | 1 | TU1e | PJ23110-sfGFP | AmpR | L1-LEU-1E, pCTK038, pCTK154 |
| pJCM543 | 1 | TU1e | PJ23111-sfGFP | AmpR | L1-LEU-1E, pCTK039, pCTK154 |
| pJCM544 | 1 | TU1e | PJ23113-sfGFP | AmpR | L1-LEU-1E, pCTK040, pCTK154 |
| pJCM545 | 1 | TU1e | PJ23115-sfGFP | AmpR | L1-LEU-1E, pCTK041, pCTK154 |
| pJCM546 | 1 | TU1e | PJ23116-sfGFP | AmpR | L1-LEU-1E, pCTK042, pCTK154 |
| pJCM547 | 1 | TU1e | PJ23117-sfGFP | AmpR | L1-LEU-1E, pCTK043, pCTK154 |
| pJCM548 | 1 | TU1e | PJ23118-sfGFP | AmpR | L1-LEU-1E, pCTK044, pCTK154 |
| pJCM549 | 1 | TU1e | PJ23119-sfGFP | AmpR | L1-LEU-1E, pCTK045, pCTK154 |
| pJCM550 | 2 | | PJ23102-sfGFP | KanR | pJCM471, pJCM534 |
| pJCM551 | 2 | | PJ23103-sfGFP | KanR | pJCM471, pJCM535 |
| pJCM552 | 2 | | PJ23104-sfGFP | KanR | pJCM471, pJCM536 |
| pJCM553 | 2 | | PJ23105-sfGFP | KanR | pJCM471, pJCM537 |
| pJCM554 | 2 | | PJ23106-sfGFP | KanR | pJCM471, pJCM538 |
| pJCM555 | 2 | | PJ23107-sfGFP | KanR | pJCM471, pJCM539 |
| pJCM556 | 2 | | PJ23108-sfGFP | KanR | pJCM471, pJCM540 |
| pJCM557 | 2 | | PJ23109-sfGFP | KanR | pJCM471, pJCM541 |
| pJCM558 | 2 | | PJ23110-sfGFP | KanR | pJCM471, pJCM542 |
| pJCM559 | 2 | | PJ23111-sfGFP | KanR | pJCM471, pJCM543 |
| pJCM560 | 2 | | PJ23113-sfGFP | KanR | pJCM471, pJCM544 |
| pJCM561 | 2 | | PJ23115-sfGFP | KanR | pJCM471, pJCM545 |
| pJCM562 | 2 | | PJ23116-sfGFP | KanR | pJCM471, pJCM546 |
| pJCM563 | 2 | | PJ23117-sfGFP | KanR | pJCM471, pJCM547 |
| pJCM564 | 2 | | PJ23118-sfGFP | KanR | pJCM471, pJCM548 |
| pJCM565 | 2 | | PJ23119-sfGFP | KanR | pJCM471, pJCM549 |

**Supplementary Table S3: Constitutive promoter expression levels.**

| Promoter | CTK Characterization (RPU) | Anderson characterization (RPU)[2]* |
|---|---|---|
| J23119 | 7.76 | |
| J23111 | 3.97 | 0.83 |
| J23104 | 1.64 | 1.03 |
| J23101 | 1.00 | 1.00 |
| J23102 | 0.98 | 1.23 |
| J23118 | 0.75 | 0.80 |
| J23107 | 0.46 | 0.51 |
| J23110 | 0.41 | 0.47 |
| J23106 | 0.37 | 0.67 |
| J23113 | 0.27 | 0.01 |
| J23108 | 0.27 | 0.73 |
| J23105 | 0.27 | 0.34 |
| J23103 | 0.27 | 0.01 |
| J23117 | 0.22 | 0.09 |
| J23109 | 0.16 | 0.06 |
| J23116 | 0.15 | 0.23 |
| J23115 | 0.08 | 0.21 |

*The RPU of the Anderson characterization was converted from normalization to J23100 to normalization to J23101.

**Supplementary Table S4: Inducible promoter dose-response parameters**

| Plasmid | Promoter | Y_min | Y_max | n | K | Dynamic range |
|---|---|---|---|---|---|---|
| pJCM435 | PTac | 2.794 | 0.014 | 1.813 | 256.617 | 194.464 |
| pJCM448 | PBAD | 1.084 | 0.015 | 2.24 | 633.467 | 73.388 |
| pJCM449 | PTet | 8.582 | 0.033 | 2.117 | 10.119 | 263.919 |

### Supplementary Table S5: NOT gate response function parameters

| Plasmid | Gate | Y_min | Y_max | n | K | Dynamic range |
|---|---|---|---|---|---|---|
| pJCM212 | A1-AmtR | 5.49 | 0.043 | 1.863 | 0.011 | 128.658 |
| pJCM213 | B1-BM3R1 | 0.479 | 0.005 | 2.378 | 0.01 | 105.956 |
| pJCM214 | B2-BM3R1 | 0.335 | 0.025 | 1.442 | 0.072 | 13.622 |
| pJCM215 | B3-BM3R1 | 0.383 | 0.094 | 0.846 | 0.843 | 4.087 |
| pJCM216 | E1-BetI | 9.191 | 0.121 | 2.081 | 0.066 | 75.669 |
| pJCM217 | F1-AmeR | 5.374 | 0.431 | 1.323 | 0.076 | 12.46 |
| pJCM218 | H1-HlyIIR | 5.049 | 0.098 | 1.7 | 0.086 | 51.392 |
| pJCM219 | I1-IcaRA | 5.029 | 0.098 | 2.575 | 0.03 | 51.362 |
| pJCM220 | L1LitR | 0.36 | 0.01 | 1.262 | 0.025 | 37.681 |
| pJCM221 | N1-LmrA | 1.211 | 0.021 | 1.429 | 0.036 | 56.789 |
| pJCM222 | P1-PhlF | 7.194 | 0.048 | 3.159 | 0.039 | 150.738 |
| pJCM223 | P2-PhlF | 7.589 | 0.05 | 2.222 | 0.103 | 152.48 |
| pJCM224 | P3-PhlF | 8.274 | 0.058 | 2.741 | 0.047 | 143.35 |
| pJCM225 | Q1-QacR | 8.525 | 0.047 | 2.534 | 0.023 | 181.924 |
| pJCM226 | Q2-QacR | 8.107 | 0.046 | 1.601 | 0.057 | 178.172 |
| pJCM227 | R1-PsrA | 7.551 | 0.173 | 1.086 | 0.073 | 43.765 |
| pJCM228 | S1-SrpR | 0.957 | 0.005 | 3.971 | 0.011 | 195.516 |
| pJCM229 | S2-SrpR | 1.837 | 0.011 | 2.558 | 0.028 | 160.211 |
| pJCM230 | S3-SrpR | 1.987 | 0.012 | 1.721 | 0.047 | 160.701 |
| pJCM231 | S4-SrpR | 1.772 | 0.01 | 2.18 | 0.051 | 170.457 |

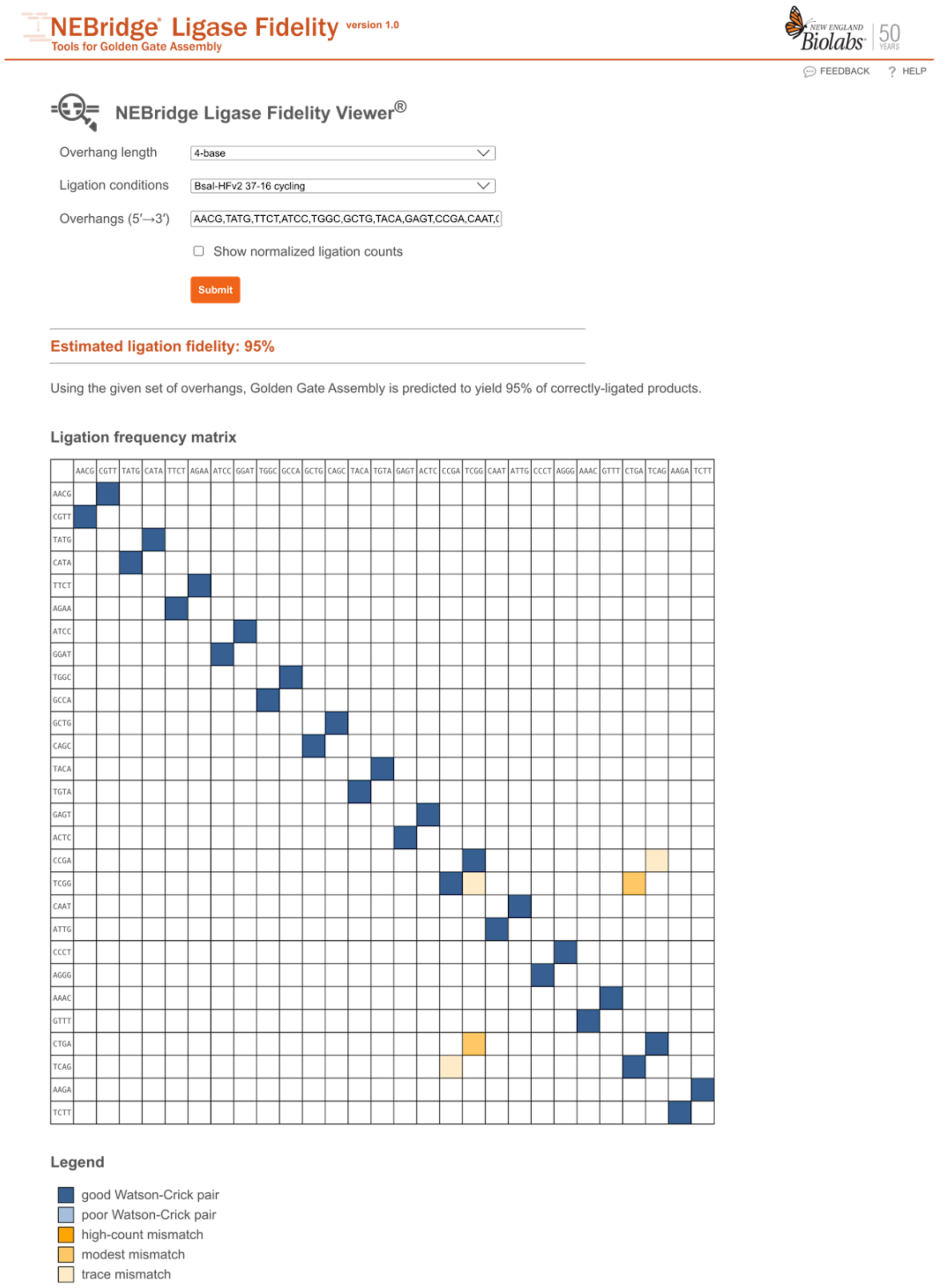


https://ligasefidelity.neb.com/viewset/run.cgi

**Supplementary Figure S1: Ligase fidelity overview of the 4-base pair overhangs employed in the CTK.** The additional overhangs, **AAAC** (between 2a and 2b), **CTGA** (between 2b and 2c), and **AAGA** (between 2c and 2d), show minimal overlap with the existing YTK overhangs. The only noticeable mismatches are between CTGA and TCGG. These are the overhangs between Type 2b and 2c and between Type 7 and 8 parts. When Type 678 backbones are used for construction of level 1 plasmids, this potential mismatch is entirely circumvented.

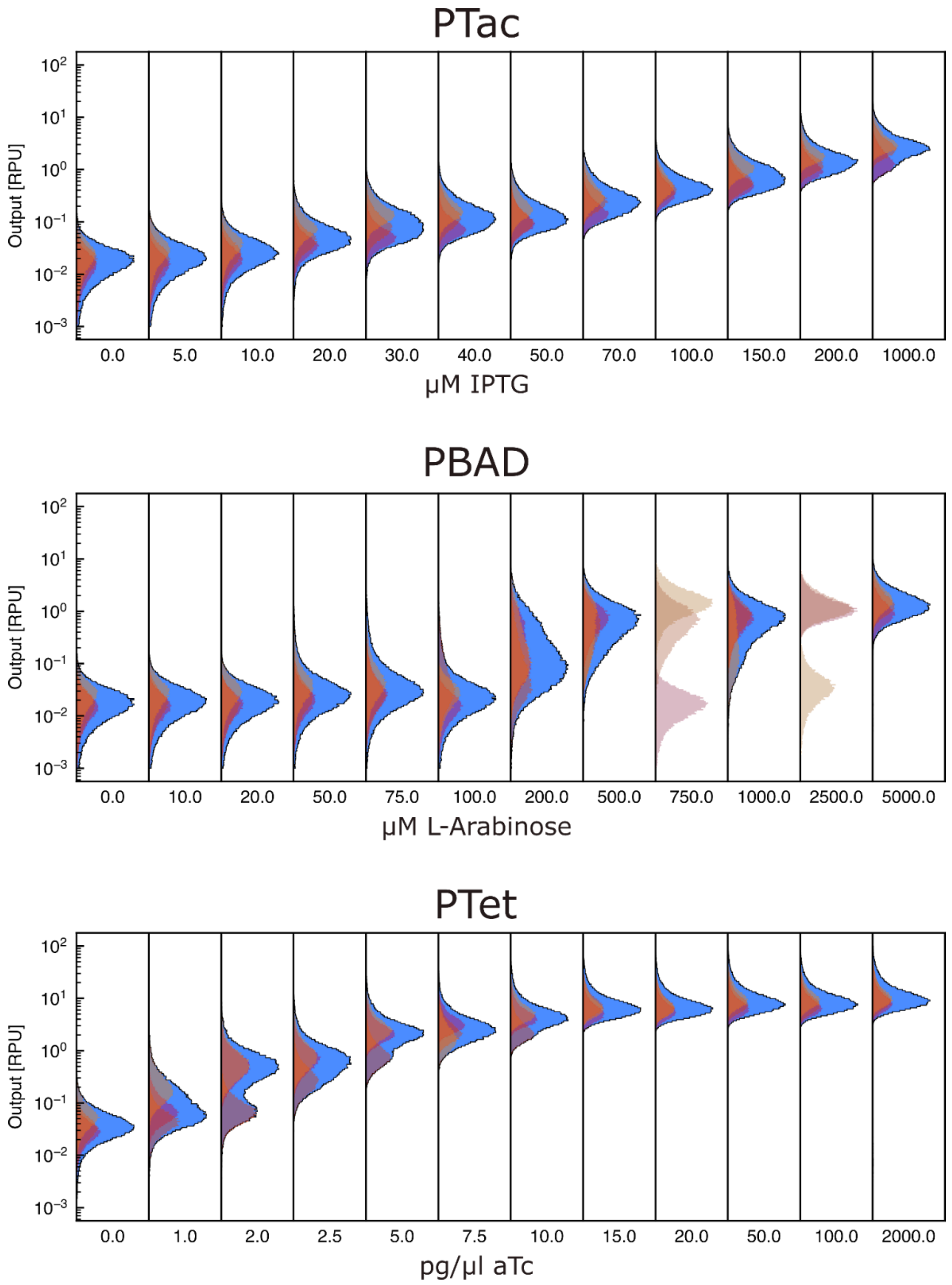


**Supplementary Figure S2: Distributions of input promoters.** The three input promoters were measured at 12 different concentrations of their respective chemical inducers. For each measurement, three replicates were performed on separate days. The individual replicates can be seen in yellow, orange and pink. The larger peak in light blue is the merged population of the replicates. Populations in light colors were outliers and not included in further calculations (see Methods). Parameters for a dose-response curve fit based on these distributions can be seen in Supplementary Table S4.

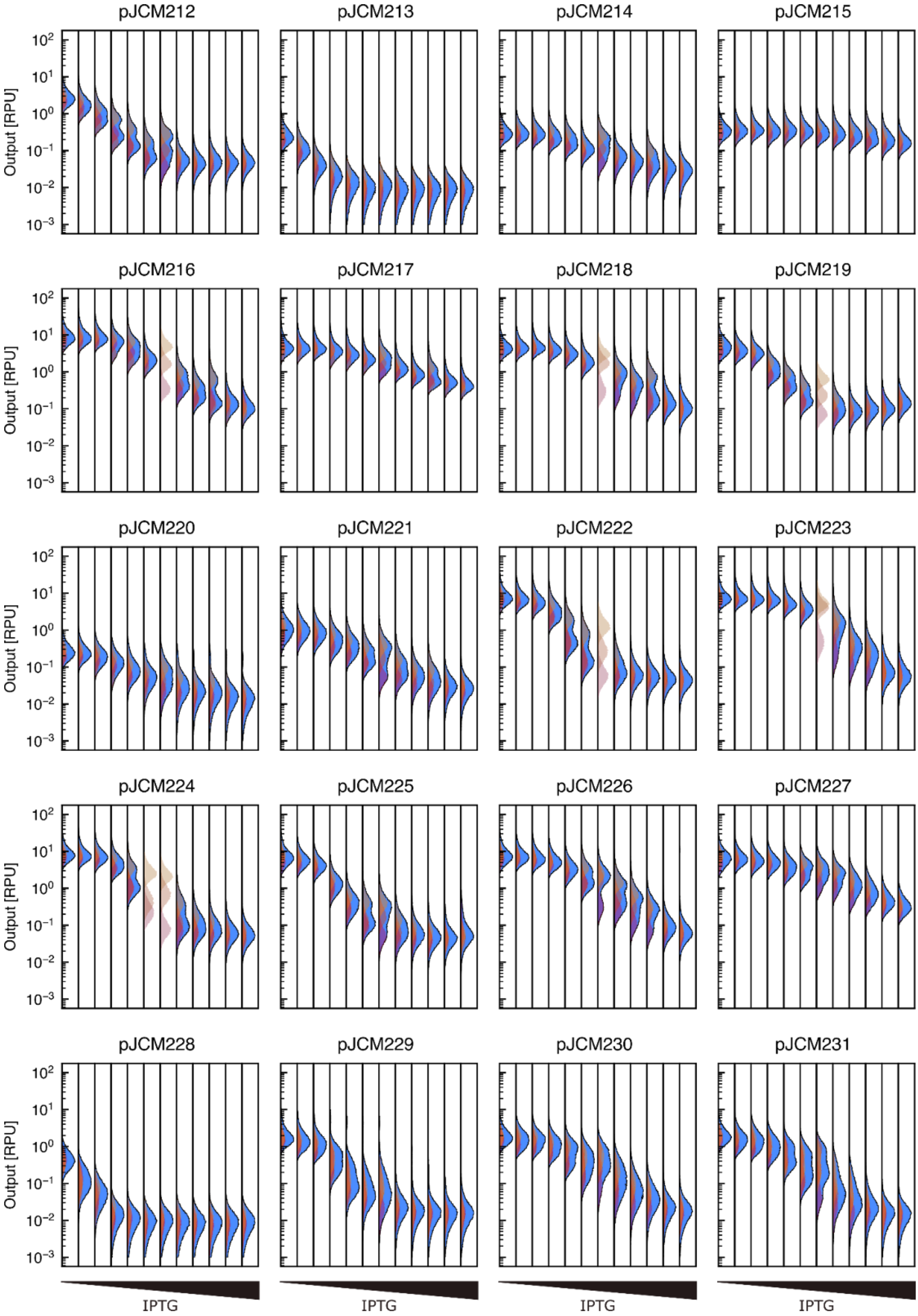


**Supplementary Figure S3: Distributions of NOT gates.** The distributions of the 20 NOT gates, each normalized to RPU. For each measurement, three replicates were performed on separate days. The individual replicates can be seen in yellow, orange and pink. The larger peak in light blue is the merged population of the replicates. Populations in light colors were outliers and not included in further calculations (see Methods). IPTG concentrations used in this experiment were (from left to right): 0, 5, 10, 20, 30, 40, 50, 70, 100, 150, 200, and 1000 μM.

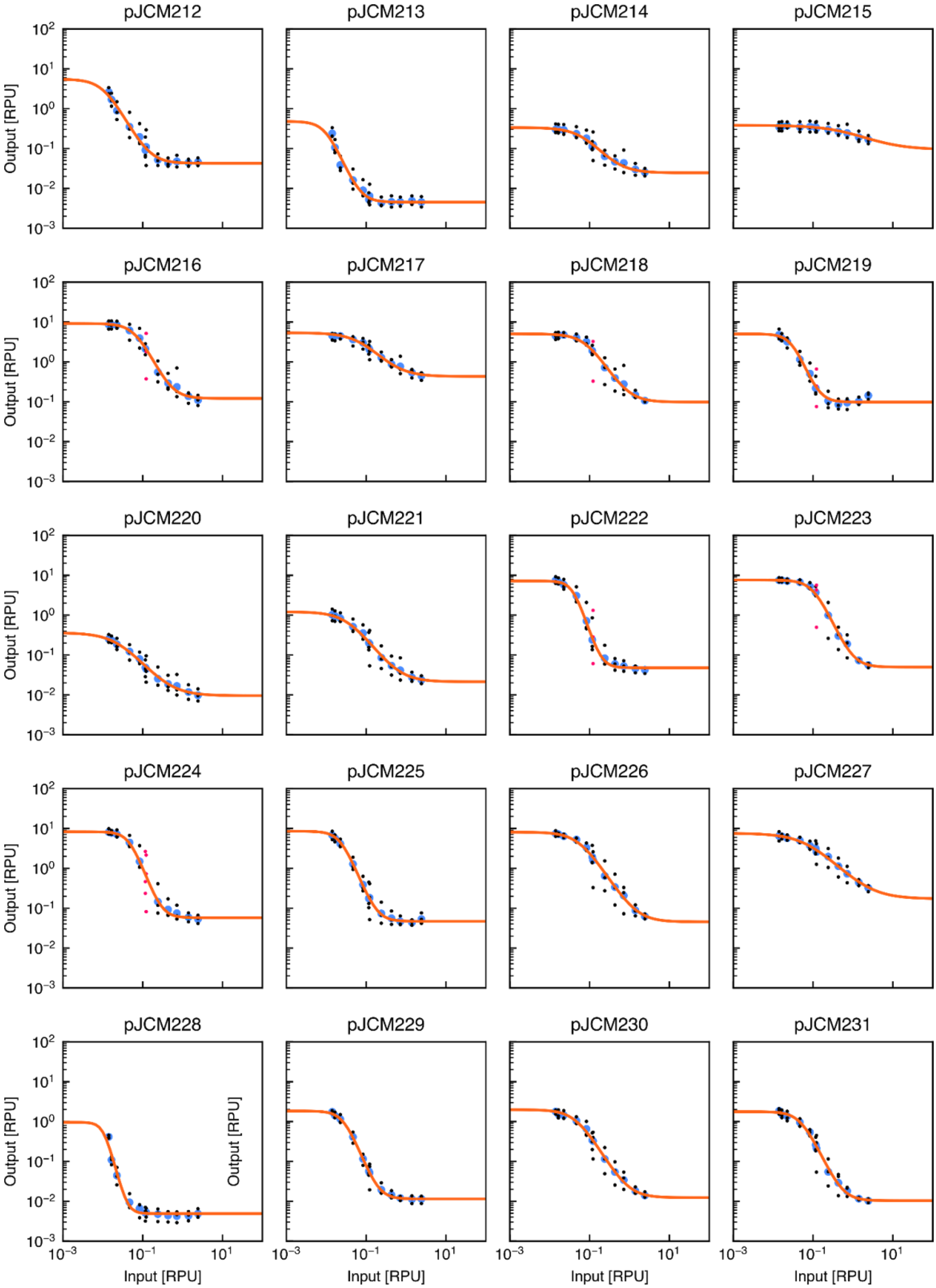


**Supplementary Figure S4: Response functions of NOT gates.** The fitted response function of the 20 NOT gates, each normalized to RPU on both input and output. The black points are individual measurements from the three replicates and the blue points are the median of the merged replicates. Points in red were outliers and not included in the merged populations. Parameters of the fitted curves can be seen in Supplementary Table S5.

**Supplementary Text: Benchmarking clustering algorithm**

For benchmarking, we used three different approaches: First, a naive approach where random nucleotides are added until the fragment is at least 300 bases long. This method will provide the baseline for comparisons between other methods, as it presents the least optimal method of packaging for *de novo* synthesis. The second approach is to randomly group fragments shorter than 300 bases together. Since this method is stochastic, the computations are repeated 10 times and both the best and the average results are reported. Third, we investigate using clustering to identify good groupings that will minimize synthesis errors while maximizing compactness.

To benchmark this tool, we chose 3 sets of DNA parts that contain many smaller parts to highlight the efficiency of the software. For all DNA sets, CDSs were excluded from this benchmarking, since they are above 300 bases and therefore would add equally to all methods tested.
The first set are the parts contained within the CTK. This set has a total of 108 parts, which breaks down into 45 promoters, 12 ribozymes, 21 RBSs, 15 CDSs, and 15 terminators.
The second set comes from the paper "Partitioning of a 2-bit hash function across 66 communicating cells" from the Voigt lab [3], was published online in the fall of 2024, and in it, the authors introduce a total of 1.1 Mb of recombinant DNA into *E. coli*. Cloning plasmids for projects of this size can be facilitated by using a hierarchical cloning scheme, such as MoClo. To get these parts into the system, many would have to be *de novo* synthesized. To create all circuits from the paper, a total of 115 parts will be needed. This breaks down into 26 promoters, 12 ribozymes, 33 RBSs, 25 CDSs, and 19 terminators.
The third set is a collection of 15 synthetic *E. coli* promoters that have a very close sequence similarity. This set was chosen as a test case to demonstrate the ability of the software to make even very similar sequences ready for synthesis.

The results of the benchmarks can be seen in the tables below. For all cases, the naive approach is by far the most expensive when compared to any other approach, and it is therefore not recommended.
When comparing the other methods described above, we see that the lowest price can be achieved by using the random clustering for two out of the three datasets. However, for the third set with the 15 promoter variants, the high sequence similarity meant that the random assignment couldn't generate a version that could be synthesized.

Using the random method for the two other DNA sets, 4 out of 10 failed for the CTK parts, and 3 out of 10 failed for the second. Here a failure is defined as having at least one sequence that is not able to be synthesized.

Using the clustering method from our tool, we were able to reduce the synthesis price to 46%, 73%, and 60%, respectively, of the naive approach for the three datasets. Furthermore, in the second set, the aggressive clustering method could decrease the synthesis price even further to 56% of the original naive approach.

**Supplementary Table S5: Benchmark of set 1, the CTK parts**

| Method | Number of fragments | Length (bp) | Price (USD) | Price compared to Naive | Success rate |
|---|---|---|---|---|---|
| Naive | 108 | 38282 | $3968.16 | 100% | 100% (1/1) |
| Aggressive clustering | 47 | 20536 | $1833.16 | 46% | 100% (1/1) |
| Clustering | 47 | 20536 | $1833.16 | 46% | 100% (1/1) |
| Average Random | 47 | 20684 | $1841.50 | 46% | 60% (6/10) |
| Best Random | 47 | 20653 | $1833.16 | 46% | |

**Supplementary Table S6: Benchmark of set 2, the parts required for the 2-bit hashing function**

| Method | Number of fragments | Length (bp) | Price (USD) | Price compared to Naive | Success rate |
|---|---|---|---|---|---|
| Naive | 115 | 46998 | 4573.10 | 100% | 100% (1/1) |
| Aggressive clustering | 57 | 31279 | 2543.10 | 56% | 100% (1/1) |
| Clustering | 63 | 33099 | 2753.10 | 60% | 100% (1/1) |
| Average Random | 55 | 30530 | 2474.43 | 54% | 70% (7/10) |
| Best Random | 55 | 30412 | 2473.10 | 54% | |

**Supplementary Table S7: Benchmark of set 3, RcsAB promoters**

| Method | Number of fragments | Length (bp) | Price (USD) | Price compared to Naive | Success rate |
|---|---|---|---|---|---|
| Naive | 15 | 4505 | 525 | 100% | 100% (1/1) |
| Aggressive clustering | - | - | - | - | 0% (0/1) |

| | | | | | |
|---|---|---|---|---|---|
| Clustering | 11 | 3320 | 385 | 73% | 100% (1/1) |
| Average Random | - | - | - | - | 0% (0/10) |
| Best Random | - | - | - | - | |
